# Excitatory drive to spinal motoneurones is necessary for serotonin to modulate motoneurone excitability via 5-HT_2_ receptors in humans

**DOI:** 10.1101/2023.04.26.538484

**Authors:** Tyler T. Henderson, Janet L. Taylor, Jacob R. Thorstensen, Justin J. Kavanagh

## Abstract

Serotonin modulates corticospinal excitability, motoneurone firing rates and contractile strength via 5-HT_2_ receptors. However, the effects of these receptors on cortical and motoneurone excitability during voluntary contractions have not been explored in humans. Therefore, the purpose of this study was to investigate how 5-HT_2_ antagonism affects corticospinal and motoneuronal excitability with and without descending drive to motoneurones. Twelve individuals (aged 24 ± 4 years old) participated in a double-blind, placebo-controlled, crossover study, whereby the 5-HT_2_ antagonist cyproheptadine was administered. Transcranial magnetic stimulation (TMS) was delivered to the motor cortex to produce motor evoked potentials (MEPs) and electrical stimulation at the cervicomedullary junction was used to generate cervicomedullary motor evoked potentials (CMEPs) in the biceps brachii at rest and during a range of submaximal elbow flexions. Evoked potentials were also obtained after a conditioning TMS pulse to produce conditioned MEPs and CMEPs (100 ms inter-stimulus interval). Compared to placebo, 5-HT_2_ antagonism reduced maximal elbow flexion torque (p = 0.004), unconditioned MEP amplitude at rest (p = 0.003), conditioned MEP amplitude at rest (p = 0.033), and conditioned MEP amplitude during contractions (p = 0.020). 5-HT_2_ antagonism also increased unconditioned CMEP amplitude during voluntary contractions (p = 0.041) but not at rest. Although 5-HT_2_ antagonism increased long-interval intracortical inhibition, net corticospinal excitability was unaffected during voluntary contractions. Given that spinal motoneurone excitability was only affected when descending drive to motoneurones was present, the current study indicates that excitatory drive is necessary for 5-HT_2_ receptors to regulate motoneurone excitability but not intracortical circuits.

**Significance statement:** Cellular and animal preparations have revealed that somatodendritic 5-HT_2_ receptors on motoneurones can modulate motoneurone excitability. However, it is mostly unknown how 5-HT_2_ receptors modulate motor cortical and motoneurone activity to generate muscle contractions in humans. Here we show that antagonism of 5-HT_2_ receptors reduced muscle responses to motor cortical stimulation only when the muscle was at rest, or when voluntary motor activity was interrupted by a conditioning TMS stimulus. In contrast, antagonism of 5-HT_2_ receptors increased the muscle response to cervicomedullary electrical stimulation, but only when descending drive to motoneurones was present. These findings not only suggest that 5-HT_2_ receptors modulate intracortical and motoneurone activity, but sustained synaptic excitation of motoneurones is required for serotonergic mechanisms to modulate motoneurones.

## INTRODUCTION

Persistent inward currents (PIC) are critical for sustained, repetitive, firing of motoneurones in mammals. Activation of PICs on the motoneurone causes significant amplification of depolarising drive to the motoneurone and evokes a strong acceleration in motoneurone firing rate (Bennett et al., 1998; Hounsgaard et al., 1988; Lee & Heckman, 1998). Monoamines such as serotonin (5-HT) and noradrenaline (NA) activate intracellular G-protein signal transduction pathways to facilitate PIC activity. Na PICs and Ca PICs amplify currents from synaptic inputs to the motoneurone up to five-fold, depending on the level of monoaminergic drive to spinal motoneurones (Heckman et al., 2009; Hultborn et al., 2003; Lee & Heckman, 2000). Brainstem monoamine pathways release 5-HT and NA onto motoneurones to regulate motor behaviour by changing motor unit recruitment and discharge characteristics. Given that synaptic current generated by descending inputs from the motor cortex are greater when more muscle force is being generated, it is also probable that monoaminergic drive is linked to contraction intensity. Although this proposition is yet to be explored in humans, feline studies have revealed that incremental increases in treadmill walking speed is near-linearly scaled to increases in activity of the 5-HT-producing raphe-nuclei in the brainstem (Jacobs & Fornal, 1997; Jacobs et al., 2002; Veasey et al., 1995).

Serotonergic modulation of motoneurone excitability is predominantly mediated via 5-HT_2_ receptors located on the soma and dendrites of motoneurones (D’Amico et al., 2013; Kavanagh & Taylor, 2022; Perrier & Cotel, 2008; Perrier & Delgado-Lezama, 2005; Perrier & Hounsgaard, 2003). Recent human studies have employed pharmacological interventions in healthy individuals to reveal that 5-HT_2_ receptor antagonism reduces maximal contraction force (Thorstensen et al., 2021, 2022), corticospinal excitability (Thorstensen et al., 2021), and motor unit discharge rates (Goodlich et al., 2022). However, each of these studies approach the motor system as a whole and do not investigate cortical and spinal contributions to muscle activation. Although the effects that 5-HT_2_ receptor antagonism have on motor cortex modulation are yet to be determined, the effect that 5-HT_2_ receptor antagonism has on spinal motoneurone excitability has been examined in a single study. Cervicomedullary electrical stimulation was employed to provide a measure of spinal motoneurone excitability to synaptic input. Interestingly, 5-HT_2_ antagonism did not affect motoneurone responsiveness to the single descending volley generated by cervicomedullary stimulation (Thorstensen et al., 2022), which was most likely due to the state of the motor system when stimulations were delivered. Given that the evoked potentials were obtained when the target muscle was at rest, it is likely that dendritic PICs were not activated. Thus, voluntary drive may be necessary to activate dendritic PICs so that overt 5-HT effects can occur at the motoneurones. This viewpoint is supported by several lines of evidence, as activation of PICs is known to require sustained synaptic excitation (Heckman et al., 2008; Heckman et al., 2003) and serotonergic effects have been most frequently observed in humans when performing strong voluntary contractions (Henderson et al., 2022; Kavanagh et al., 2019).

The purpose of this study was to investigate how 5-HT_2_ receptor antagonism affects corticospinal and motoneuronal excitability. To achieve this, transcranial magnetic stimulation (TMS) was delivered to the motor cortex to produce motor evoked potentials (MEPs), and electrical cervicomedullary stimulation was delivered to produce cervicomedullary motor evoked potentials (CMEPs). Evoked potentials were measured in the biceps brachii at rest and during submaximal elbow flexions. Given that MEPs and CMEPs are heavily influenced by descending drive to the muscle, MEPs and CMEPs were also obtained during a period of myoelectric silence that was induced by a preceding TMS pulse. This conditioning stimulus enabled the assessment of intracortical excitability during a transient inhibition of corticospinal output and the assessment of spinal motoneurone excitability during a period of disfacilitation (McNeil et al., 2013; McNeil et al., 2009). Unconditioned and conditioned MEPs and CMEPs were assessed before, and after, the administration of the 5-HT_2_ antagonist cyproheptadine. Since 5-HT_2_ antagonism can increase measures of motor cortical inhibition (Thorstensen et al., 2021) and reduce estimates of PICs at motoneurones (D’Amico et al., 2013; Goodlich et al., 2023), we hypothesised that 5-HT_2_ receptor antagonism would increase intracortical inhibition, and reduce motoneurone excitability during voluntary contractions.

## METHODS

### Experimental design

This study was a human, double-blind, placebo-controlled, two-way cross-over trial. Participants attended the laboratory on two occasions where a placebo or cyproheptadine capsule was ingested. The session that each drug was administered was counterbalanced to avoid order effects. Testing sessions were separated by a minimum of 7 days to ensure residual drug effects did not carry over into the second session. Testing procedures were performed pre-pill ingestion (baseline) and post-pill ingestion for both sessions.

### Participants and ethical approval

Fifteen healthy individuals participated in this study (24 ± 4 yr, 4 female). Each participant was screened using a medical history questionnaire which contained exclusion criteria specific to upper limb musculoskeletal injury, cyproheptadine contraindications, magnetic stimulation, and electrical stimulation. Recruited individuals also attended a familiarisation session to ensure that participants could tolerate stimulation, and to confirm that it was possible to evoke an electromyographic (EMG) response in the target muscle with motor cortical stimulation and cervicomedullary stimulation. Following screening, twelve participants (24 ± 4 yr, 2 female) continued to participate in experimental testing sessions. Participants were asked to refrain from any stimulants and exercise for 18 hr prior to testing sessions. Ethical approval was obtained via the Human Research Ethics committee at Griffith University (Griffith University Reference Number: 2020/264), and written informed consent was obtained prior to testing. All testing procedures conformed to the standards set by the *Declaration of Helsinki* except for registration in a database.

### Drug intervention

A single 8 mg oral dose of the 5-HT_2_ receptor antagonist cyproheptadine, or a placebo, was administered in opaque capsules in separate testing sessions. Cyproheptadine is a competitive antagonist of the 5-HT_2_ receptor and has a high binding affinity for 5-HT_2A_, 5-HT_2B_, and 5-HT_2C_ receptor subtypes (Honrubia et al., 1997). Post-pill testing commenced 2.5 hr after drug administration to coincide with previously reported drug effects that cyproheptadine has on reflex responses and motoneurone activity (D’Amico et al., 2013; Goodlich et al., 2022; Thorstensen et al., 2021, 2022; Wei et al., 2014).

### Participant setup and electromyography

Participants were seated with their right arm secured firmly in a custom-designed transducer with a non-compliant strap. The transducer placed the elbow and shoulder in 90 degrees of flexion, where a load cell (200 kg capacity, PT4000 S-Type, PT Ltd., NZ) was calibrated to measure isometric elbow flexion torque. A computer monitor was positioned ∼1 m in front of participants at eye level to provide real-time feedback on torque generation. Bipolar surface EMG signals were obtained from the right biceps brachii of each participant via Ag/AgCl electrodes (Kendall ARBO, 24 mm diameter). One electrode was placed over the mid-point of the muscle belly and one electrode over the distal tendon of the biceps brachii. EMG was amplified (x 100), and bandpass filtered between 10 and 1000 Hz using a 2nd order Butterworth filter (CED 1902, Cambridge Electronic Design Ltd., UK). Torque and EMG data were sampled at 2000 Hz using a Power 1401 data acquisition interface with Signal software (version 6, Cambridge Electronic Design Ltd., UK).

### Brachial plexus stimulation

To evoke a maximal compound muscle action potential (M_MAX_) in the biceps brachii, a constant current stimulator was used to deliver single electrical stimuli to nerve fibres of the brachial plexus (0.2 ms pulse width, DS7AH, Digitimer Ltd., UK). A surface cathode was positioned over the right supraclavicular fossa (Erb’s point) and a surface anode placed on the right acromion process. Stimulator intensity was set at 130% of the intensity required to produce M_MAX_ in the EMG signal of the resting biceps brachii (average intensity across both testing sessions: 135 ± 46 mA). Stimulator intensity remained fixed throughout the testing sessions and was not adjusted after pill ingestion.

### Motor cortical stimulation

Two Magstim 200^2^ stimulators (connected via a BiStim module, Magstim Co, Dyfed, UK) were used to deliver magnetic stimulation to the motor cortex via a circular coil (90 mm diameter, Magstim Co., UK) positioned over the vertex. Three different forms of stimuli were used to evoke responses in the right biceps brachii: a single conditioning stimulus, a single test stimulus (unconditioned response), and a test stimulus delivered 100 ms after a conditioning stimulus (conditioned response) (McNeil et al., 2009). The test stimulus intensity was set to produce a motor evoked potential (MEP) with an amplitude ∼60% of peak-to-peak amplitude of M_MAX_ in biceps brachii during elbow flexions performed at 20% MVC (70 ± 15% maximum stimulator output, MSO). The conditioning stimulus intensity was set to elicit a silent period greater than 200 ms during elbow flexions performed at 50% MVC (76 ± 13% MSO). Once established, the stimulation intensities remained fixed throughout the testing session. Stimulation intensities were set prior to pill ingestion to account for the potential lengthening of the TMS-induced silent period after cyproheptadine ingestion which would indicate stronger inhibitory effects on the motor cortex for the drug condition (Thorstensen et al., 2021).

### Cervicomedullary stimulation

A constant current stimulator (0.2 ms pulse width, DS7AH, Digitimer Ltd., UK) was used to deliver electrical stimuli to the cervicomedullary junction to elicit cervicomedullary motor evoked potentials (CMEPs) in the right biceps brachii. CMEP responses were evoked with single cervicomedullary stimuli (unconditioned CMEP), as well as 100 ms after a conditioning TMS pulse (conditioned CMEP). A surface cathode and anode were placed slightly (∼ 1 cm) inferior and medial to the left and right mastoid process. The stimulus intensity was set to produce a CMEP with a similar amplitude to the test MEP (∼ 60% of peak-to-peak amplitude of M_MAX_ during elbow flexions performed a 20% MVC, 186 ± 24 mA). These stimulator intensities remained fixed throughout the testing sessions and were not adjusted after pill ingestion. To ensure descending corticospinal axons were activated, and not cervical nerve roots, the latency of CMEP onset (∼8 ms) was monitored throughout testing. Most notably, a sudden ∼2 ms decrease in latency suggests that cervical axons are activated (Petersen et al., 2002; Taylor, 2006; Taylor & Gandevia, 2004; Ugawa et al., 1991). CMEP measurements were not obtained from one participant (female) as there was a sudden shift in CMEP latency (∼ 6 ms) with an increase in stimulation intensity.

### Experiment protocols

#### MVC torque

Participants performed 4 brief maximal contractions (∼ 4 s) to establish maximal torque output. The greatest torque generated during these contractions was deemed the participant’s MVC, and submaximal torque targets were calculated from this value. TMS and electrical stimulator intensities were determined from 20% and 50% MVCs at this point in the experiment (see *Motor cortical stimulation* and *Cervicomedullary stimulation* sections).

#### Pre-pill (baseline) protocol

Brachial plexus stimulation, motor cortical stimulation, and cervicomedullary stimulation were used to evoke M_MAX_, unconditioned MEP, conditioned MEP, unconditioned CMEP, and conditioned CMEP responses. Evoked potentials were measured from participants at rest as well as during brief (∼ 5 s) elbow flexions of 20% MVC, 50% MVC and 80% MVC. A single evoked response was obtained from a single contraction, where each stimulus was delivered ∼ 3 s into each contraction. Four evoked responses were obtained for each stimulation type at each contraction intensity. The order of contraction intensity was randomised with sufficient rest periods (1-4 min) provided between contractions to minimise fatigue.

#### Post-pill protocol

Participants performed 4 brief maximal contractions (∼ 4 s) to establish post-pill MVC. However, submaximal torque targets were based on pre-pill MVC. This ensured that the effect of 5-HT_2_ receptor antagonism could be examined with respect to each submaximal contraction intensity rather than with respect to new submaximal contraction intensities created by drug-effects in post-pill MVCs. An additional four 50% MVCs were performed post-pill, where a single TMS conditioning stimulus was delivered to ensure that the TMS-evoked silent period remained greater than 200 ms post-pill. Like the pre-pill protocol, the five types of evoked potential measurements were once again obtained from participants at rest and during each contraction intensity. Four trials were performed for each stimulation type, at each contraction intensity. The order of contraction intensity remained the same as the pre-pill protocol.

### Data analysis

All measures were calculated offline using Signal software (version 7, Cambridge Electronic Design). Root mean squared (RMS) amplitude of voluntary EMG was calculated from a 200 ms window immediately prior to stimulations. Silent period duration was the time from the onset of the stimulus (i.e., single conditioning TMS, single test TMS and single cervicomedullary stimulation) to the return of voluntary EMG (as determined by visual inspection). The peak-to-peak amplitude of M_MAX_, as well as conditioned and unconditioned MEP and CMEP responses were calculated for each contraction that was performed. For all variables, a window for the amplitude calculation was defined from the initial deflection in the EMG signal from baseline to the second crossing of the 0-mV line. Evoked potentials were normalised to the average M_MAX_ obtained for the same contraction intensity within each testing session and within each testing day. The percentage change (Δ) in evoked potentials within each testing session was determined by using the formula: 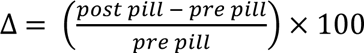, so that changes are presented as percentage difference from pre-pill (baseline). A positive change score reflects an increase in the magnitude of the post-pill measurement compared to the pre-pill measurement.

### Statistical analysis

Paired t-tests were used to determine if pre-pill MVC torque differed between testing sessions. Paired t-tests were also used to examine if post-pill changes in MVC torque, and post-pill changes in evoked potentials, differed between the cyproheptadine and placebo sessions. Mauchly’s test of sphericity was applied to all measures obtained during submaximal voluntary contractions, and Greenhouse-Geisser corrections were applied to non-spherical data. Two-way repeated measures ANOVA were used to examine the main effect of conditioning stimulus (unconditioned vs conditioned), main effect of contraction intensity (20% MVC, 50% MVC, 70% MVC), and conditioning stimulus by contraction intensity interaction, for MEP and CMEP responses. Two-way repeated measures ANOVA were also used to examine the main effect of drug (cyproheptadine vs placebo), contraction intensity (20% MVC, 50% MVC, 70% MVC), and drug by contraction intensity interaction, for post-pill changes in MEP, CMEP, RMS EMG, and M_MAX_ responses, as well as silent period durations. Tukey’s multiple comparison tests were applied to data when significant interaction effects were detected. All statistical tests were performed using SPSS Statistics (v.25, IBM Corp). Statistical significance was set at p < 0.05 and effect sizes are reported as partial eta squared (η^2^). All data are presented as mean ± standard deviation in tables and figures.

## RESULTS

### Maximal torque generation

MVC torque was measured pre- and post-pill ingestion for placebo and cyproheptadine testing conditions. Pre-pill measures of maximal torque were not significantly different between testing days (t = −1.274, p = 0.229, Figure 1A), which suggests that participants were able to generate similar elbow flexion forces for each testing session before any intervention occurred. To assess the effect that cyproheptadine had on maximal contractions, the change in force was calculated for MVC torque within each testing session (post-pill normalised to pre-pill torque), where cyproheptadine caused a significantly greater reduction in MVC torque compared to placebo (t = 3.645, p = 0.003, Figure 1B).

**Figure 1.**
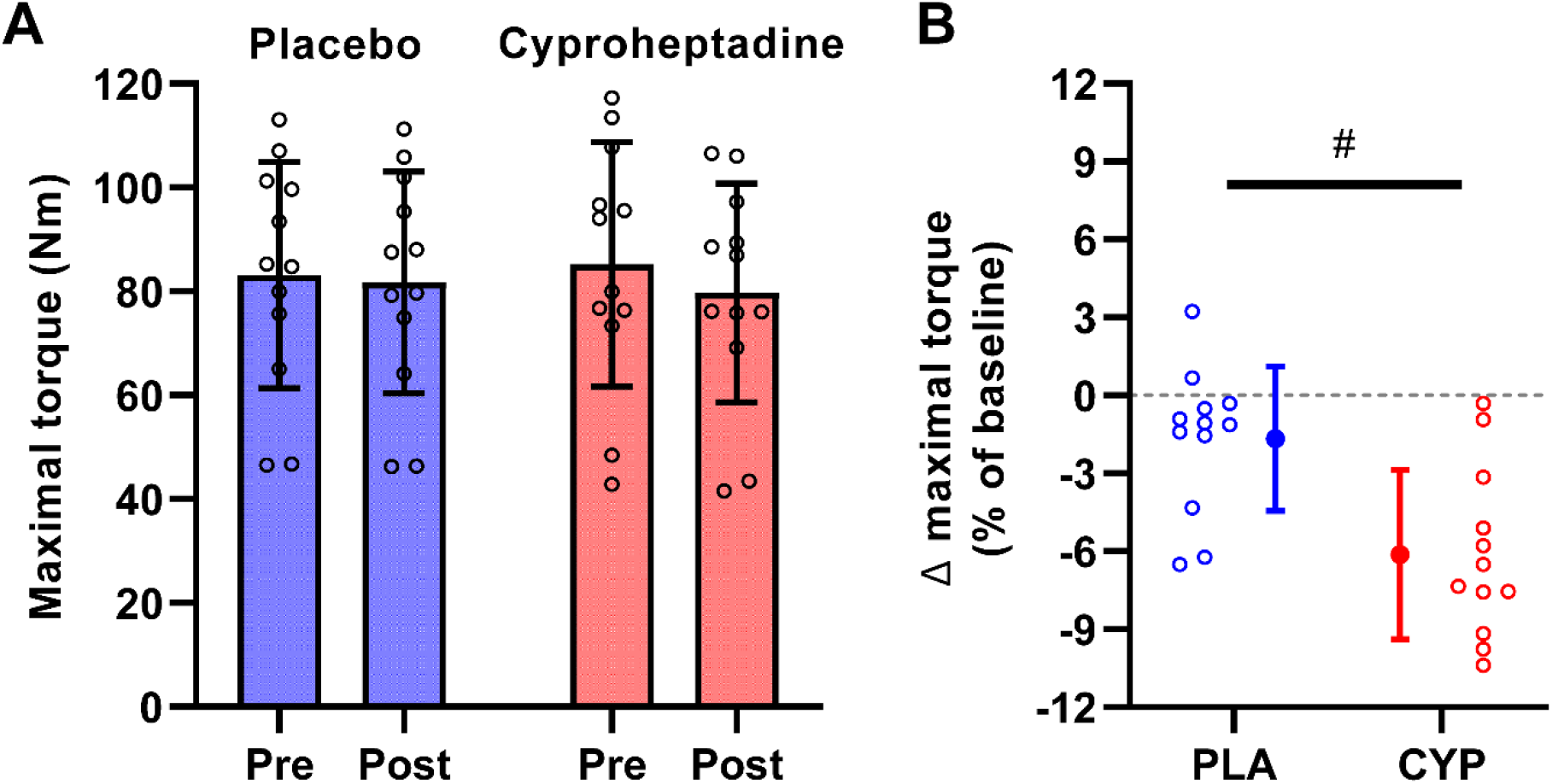
MVC torque during maximal elbow flexions. MVCs were measured pre-pill ingestion and post-pill ingestion for both the placebo session and the cyproheptadine session (A). Post-pill ingestion values were normalised to pre-pill ingestion values so that MVC is presented as a percentage change from pre-pill ingestion values (B). Data are presented as the group mean (n = 12, 2 female) and standard deviation for the placebo session (blue, PLA) and cyproheptadine session (red, CYP). Individual data are presented as circles. The hash symbol indicates a significant difference between placebo and cyproheptadine (p < 0.05).

### Unconditioned and conditioned evoked potentials during submaximal contractions

Representative data are presented for a single participant in Figure 2. The amplitudes of unconditioned MEPs measured from the biceps brachii were low when the elbow flexors were at rest (Figure 2A). However, the performance of voluntary elbow flexions produced clear MEPs in the biceps brachii. With respect to the conditioned MEPs, delivery of a TMS conditioning stimulus prior to the test stimulus resulted in a barely discernible MEP with the muscle at rest, which progressively increased to 50% MVC, and then remaining at a similar amplitude at 80% MVC. Unconditioned CMEP amplitude progressively increased from rest to 50% MVC, and then remained at a similar amplitude at 80% MVC (Figure 2B). The delivery of a TMS conditioning pulse prior to the test stimulus produced a clear CMEP in the resting biceps brachii. The amplitude of conditioned CMEPs progressively decreased from rest, to 20% MVC, to 50% MVC, to 80% MVC.

**Figure 2.**
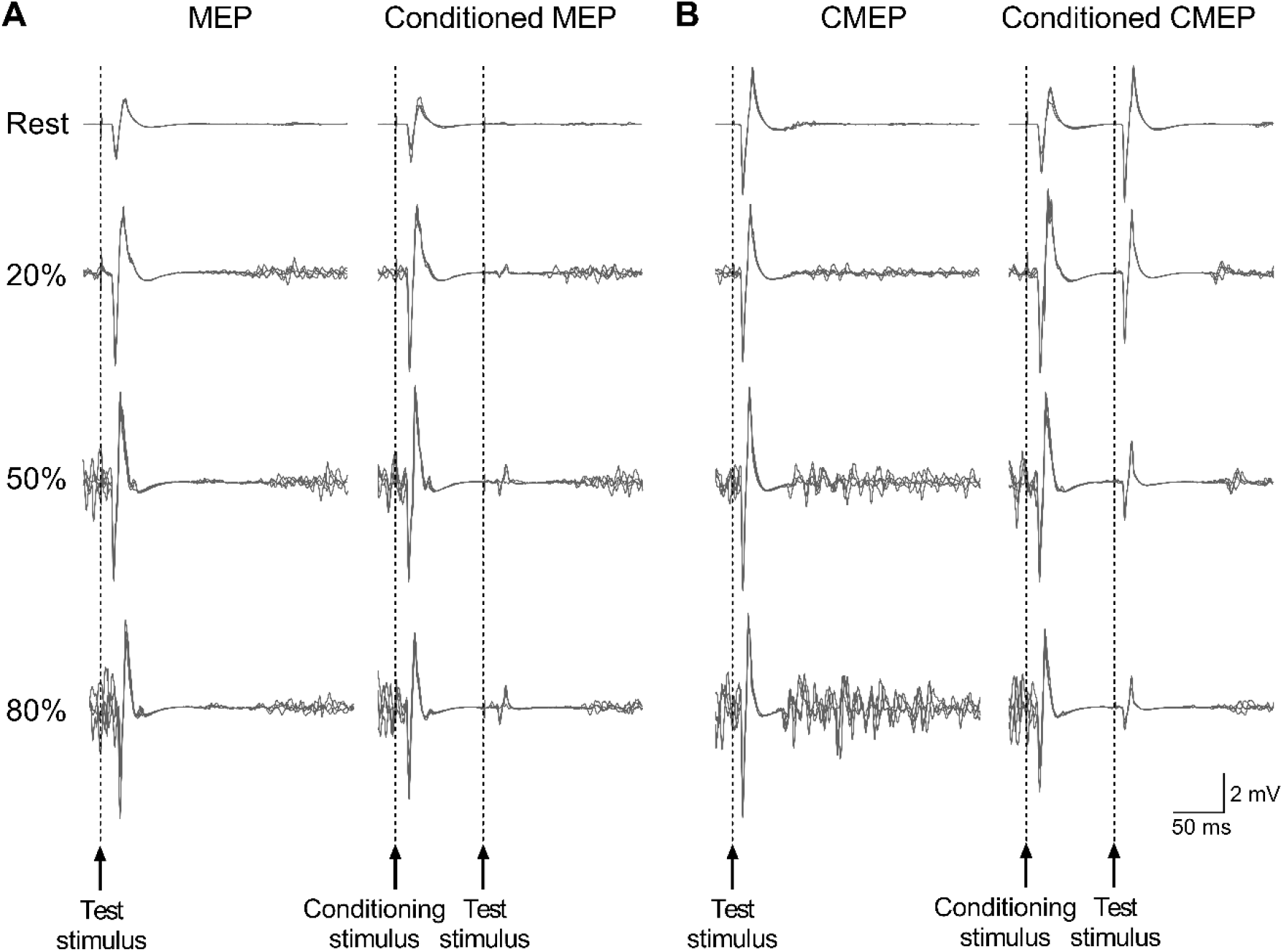
Representative evoked potentials for a single participant. MEPs were produced with a single TMS pulse to the motor cortex with, and without, a TMS conditioning pulse that was delivered 100 ms prior to the test pulse (A). CMEPs were produced with a single electrical pulse delivered at the cervicomedullary junction with, and without, a TMS conditioning pulse that was delivered 100 ms prior to the test pulse (B). All responses were obtained from biceps brachii EMG during rest and during elbow flexions at 20%, 50% and 80% of maximal torque.

As there were no significant differences for pre-pill measures between the placebo and cyproheptadine sessions (see Table 1), pre-pill evoked responses were pooled for the purpose of characterising MEPs and CMEPs without the influence of an intervention (Figure 3). When the elbow flexors were at rest there were no differences between unconditioned and conditioned responses for MEP amplitude (t = 1.052, p = 0.315, Figure 3A) and CMEP amplitude (t = 0.688, p = 0.507, Figure 3C). However, during voluntary elbow flexions, significant differences were identified between unconditioned and conditioned MEPs. There were main effects of conditioning stimulus (F(1, 11) = 10.389, p = 0.008, η^2^ = 0.486), contraction intensity (F(2, 22) = 27.774, p < 0.001, η^2^ = 0.716), and a conditioning stimulus by contraction intensity interaction (F(2, 22) = 21.562, p < 0.001, η^2^ = 0.662) detected for MEP amplitude. Post hoc analysis revealed that the conditioned MEP amplitude was lower than the unconditioned MEP amplitude during 20% (p < 0.001) and 50% (p = 0.022) contractions but not during 80% (p = 0.291) contractions. Significant differences were also identified between unconditioned and conditioned CMEPs during voluntary contractions. There were main effects of conditioning stimulus (F(1, 10) = 621.724, p < 0.001, η^2^ = 0.984), contraction intensity (F(2, 20) = 11.219, p < 0.001, η^2^ = 0.529), and a conditioning stimulus by contraction intensity interaction (F(2, 20) = 45.190, p < 0.001, η^2^ = 0.819) detected for CMEP amplitude. Post hoc analysis revealed that the conditioned CMEP amplitude was reduced during all contraction intensities (all p < 0.001).

**Figure 3.**
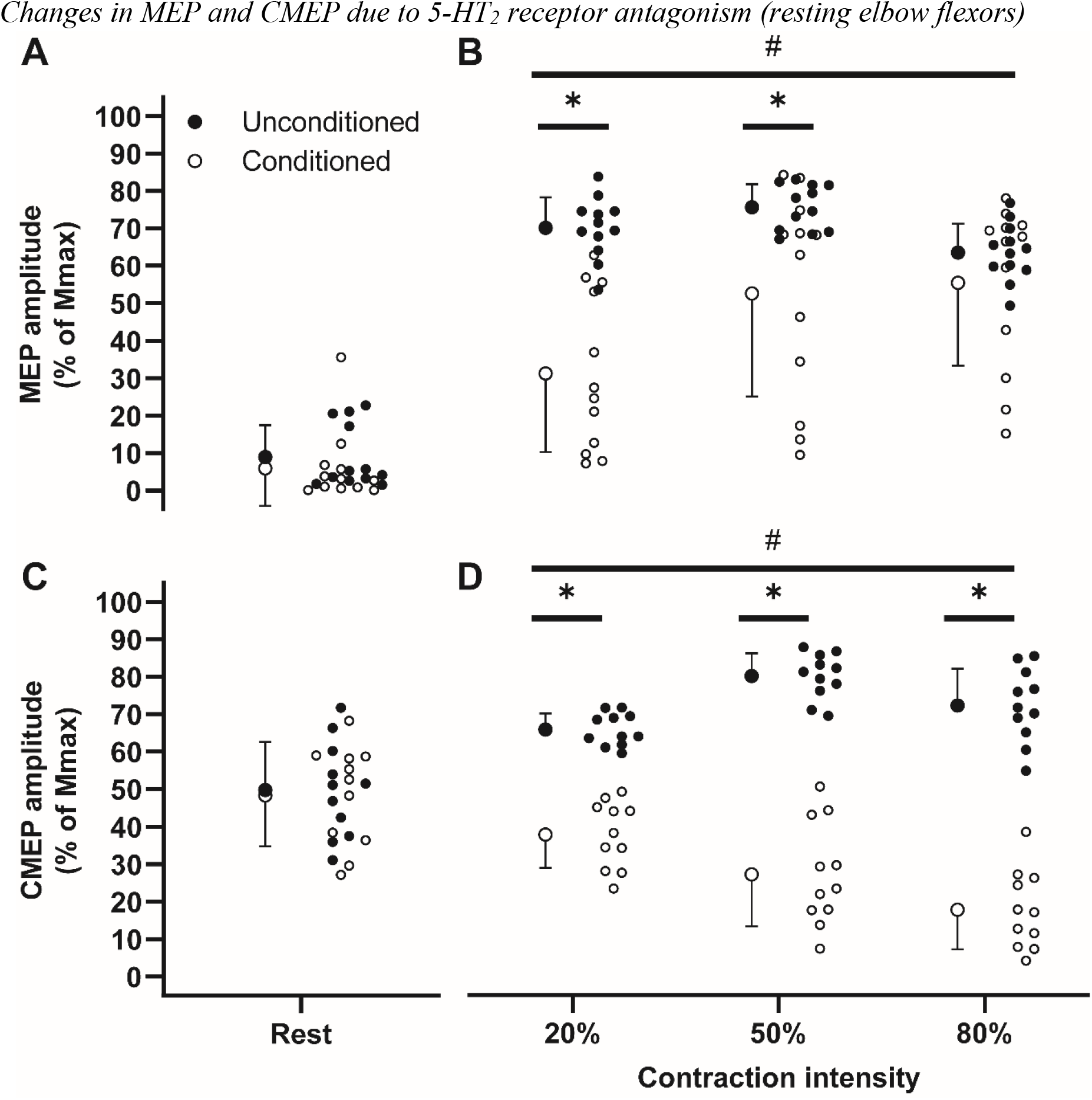
Evoked potentials obtained from healthy participants prior to intervention. Unconditioned and conditioned responses were measured from the biceps brachii of 12 healthy individuals (MEPs: n = 12, CMEPs: n = 11). MEPs were obtained with the muscle at rest (A) and during submaximal contractions (B). CMEPs were also obtained with the muscle at rest (C) and during submaximal contractions (D). Contraction intensity was set to 20% MVC, 50% MVC, and 80% MVC. Data are pooled from pre-pill measurements across testing two sessions and are presented as the mean and standard deviation of the mean. Hash symbols indicate significant main effect of contraction intensity (p < 0.05) and asterisks represent a significant post hoc comparison between conditioned and unconditioned responses (p < 0.05).

**Table 1.** Table of pre-pill ingestion measures for placebo and cyproheptadine testing sessions.

|  |  | Unconditioned<br>MEP amplitude<br>(% M<sub>MAX</sub>) | Conditioned MEP<br>amplitude<br>(% M<sub>MAX</sub>) | Unconditioned<br>CMEP amplitude<br>(% M<sub>MAX</sub>) | Conditioned CMEP<br>amplitude<br>(% M<sub>MAX</sub>) |
| --- | --- | --- | --- | --- | --- |
| Rest | PLA | 8.51 ± 8.50 | 6.09 ± 10.95 | 52.99 ± 12.81 | 47.58 ± 14.92 |
| Rest | CYP | 9.66 ± 8.81 | 5.99 ± 9.25 | 46.61 ± 16.52 | 48.95 ± 16.62 |
| *Paired-samples*<br>*T-test* | T-statistic (p-value) | -1.230 (0.241) | 0.088 (0.931) | 1.443 (0.180) | -0.278 (0.787) |
| 20% | PLA | 63.25 ± 10.14 | 25.40 ± 22.75 | 67.63 ± 6.44 | 39.21 ± 9.98 |
| 20% | CYP | 63.90 ± 7.00 | 24.12 ± 20.50 | 64.14 ± 3.56 | 36.58 ± 11.24 |
| 50% | PLA | 69.08 ± 6.20 | 46.73 ± 30.11 | 81.04 ± 6.60 | 28.03 ± 15.53 |
| 50% | CYP | 69.18 ± 7.67 | 45.48 ± 25.35 | 79.30 ± 9.96 | 26.42 ± 13.09 |
| 80% | PLA | 54.40 ± 7.28 | 47.35 ± 25.50 | 74.48 ± 9.40 | 18.63 ± 10.28 |
| 80% | CYP | 59.67 ± 10.18 | 50.50 ± 19.98 | 70.22 ± 13.27 | 16.85 ± 11.20 |
| *Two-way repeated measures*<br>*ANOVA* |  |  |  |  |  |
| Testing day | F-statistic<br>(p-value) | 1.595 (0.233) | 0.006 (0.938) | 4.598 (0.058) | 1.164 (0.306) |
| Intensity | F-statistic<br>(p-value) | **21.115 (< 0.001)\*** | **24.615 (< 0.001)\*** | **19.926 (< 0.001)\*** | **28.195 (< 0.001)\*** |
| Interaction | F-statistic<br>(p-value) | 2.696 (0.090) | 1.492 (0.247) | 0.211 (0.812) | 0.062 (0.940) |

Mean and standard deviations are reported. Paired-samples T-tests were used to determine if evoked responses obtained at rest prior to placebo and cyproheptadine ingestion were different between testing days. Two-way repeated measures ANOVA was used to determine any differences in evoked responses prior to placebo or cyproheptadine ingestion during voluntary contractions. Asterisks (\*) reflect significant main effects. Significance was set at $p < 0.05$ . MEP = motor evoked potential, M<sub>MAX</sub> = maximal compound muscle action potential, CMEP = cervicomedullary motor evoked potential, ms = milliseconds, PLA = placebo, CYP = cyproheptadine.

### Changes in MEP and CMEP due to 5-HT_2_ receptor antagonism (resting elbow flexors)

Unconditioned and conditioned responses were obtained pre- and post-pill ingestion while the elbow flexors were at rest. Post-pill ingestion values were normalised to pre-pill ingestion values so that MEPs and CMEPs are presented as a percentage change from pre-pill ingestion. Following pill ingestion, cyproheptadine caused a significantly greater reduction in unconditioned MEP amplitude (t = 3.812, p = 0.003) and conditioned MEP amplitude (t = 2.431, p = 0.033) compared to placebo. In contrast, there were no significant differences between cyproheptadine and placebo for unconditioned CMEP amplitude (t = −1.012, p = 0.335) or conditioned CMEP amplitude (t = 2.197, p = 0.053). M_MAX_ was similar between the placebo and cyproheptadine sessions which suggests that drug-effects were associated with the CNS rather than the muscle (Table 2).

**Table 2.** RMS EMG and maximal M-wave amplitudes across all contraction intensities.

| Intensity | Session | RMS EMG amplitude |  | M<sub>MAX</sub> amplitude |  |
| --- | --- | --- | --- | --- | --- |
|  |  | Pre-pill ingestion<br>(% MVC amp.) | Change from<br>pre-pill<br>(% of baseline) | Pre-pill ingestion<br>(mV) | Change from<br>pre-pill<br>(% of baseline) |
| MVC | PLA | 0.91 ± 0.29 | 2.11 ± 9.16 | - | - |
| MVC | CYP | 0.89 ± 0.25 | -3.18 ± 7.66 | - | - |
| Rest | PLA | - | - | 8.46 ± 2.64 | 6.09 ± 7.76 |
| Rest | CYP | - | - | 8.28 ± 2.82 | 3.93 ± 7.79 |
| *Paired samples T-test* |  |  |  |  |  |
| Testing day (pre-pill) or<br>drug (change scores) | T-statistic (p-value) | 0.675 (0.513) | 1.797 (0.099) | 0.672 (0.515) | 0.728 (0.481) |
| 20% | PLA | 10.58 ± 3.24 | 3.84 ± 22.40 | 9.11 ± 2.28 | 4.09 ± 7.71 |
| 20% | CYP | 10.80 ± 3.66 | 2.20 ± 23.48 | 9.19 ± 2.57 | -0.19 ± 6.68 |
| 50% | PLA | 38.71 ± 12.12 | 3.03 ± 18.43 | 9.07 ± 2.43 | 3.74 ± 7.78 |
| 50% | CYP | 35.85 ± 6.92 | 5.71 ± 18.26 | 9.35 ± 2.40 | -0.56 ± 5.63 |
| 80% | PLA | 63.06 ± 20.82 | 8.89 ± 18.58 | 9.12 ± 2.11 | 0.17 ± 9.05 |
| 80% | CYP | 64.16 ± 12.39 | 4.29 ± 15.24 | 9.31 ± 2.15 | -2.12 ± 7.04 |
| *Two-way repeated measures ANOVA* |  |  |  |  |  |
| Testing day (pre-pill) or<br>drug (change scores) | F-statistic (p-value) | 0.028 (0.871) | 0.031 (0.863) | 0.273 (0.612) | 2.311 (0.157) |
| Intensity | F-statistic (p-value) | 186.338 (< 0.001)* | 0.261 (0.773) | 0.050 (0.952) | 1.649 (0.215) |
| Interaction | F-statistic (p-value) | 0.394 (0.679) | 0.315 (0.733) | 0.537 (0.592) | 0.238 (0.791) |

Paired-samples T-tests were used to determine if measures obtained at rest or during maximal contractions (MVC) were different between testing days. Two-way repeated measures ANOVA was used to determine if differences occurred in measures prior to placebo or cyproheptadine ingestion, or differences in calculated change scores during voluntary contractions. Asterisks (\*) reflect significant main effects. Significance was set at $p < 0.05$ .

### Changes in MEP and CMEP due to 5-HT_2_ receptor antagonism (submaximal elbow flexions)

There were no significant changes in unconditioned MEP amplitude due to cyproheptadine or the submaximal contraction being performed, as there was no main effect of drug (p = 0.157), contraction intensity (p = 0.195), or drug by contraction intensity interaction (p = 0.113) detected for unconditioned MEP change (Figure 5A). However, drug-effects were detected when the MEP was preceded by a TMS conditioning stimulus, as cyproheptadine caused a significantly greater reduction in conditioned MEP amplitude compared to placebo (main effect of drug: F(1, 11) = 7.461, p = 0.020, η^2^ = 0.404, Figure 5B). Changes in the conditioned MEP were not dependent on the intensity of the elbow flexion, as there was no main effect of contraction intensity (p = 0.657), and no drug by contraction intensity interaction (p = 0.944) identified for conditioned MEP amplitude. There was a significant increase in the TMS induced silent period following cyproheptadine ingestion compared to placebo for single conditioning and test stimuli (see Table 3).

**Figure 4.**
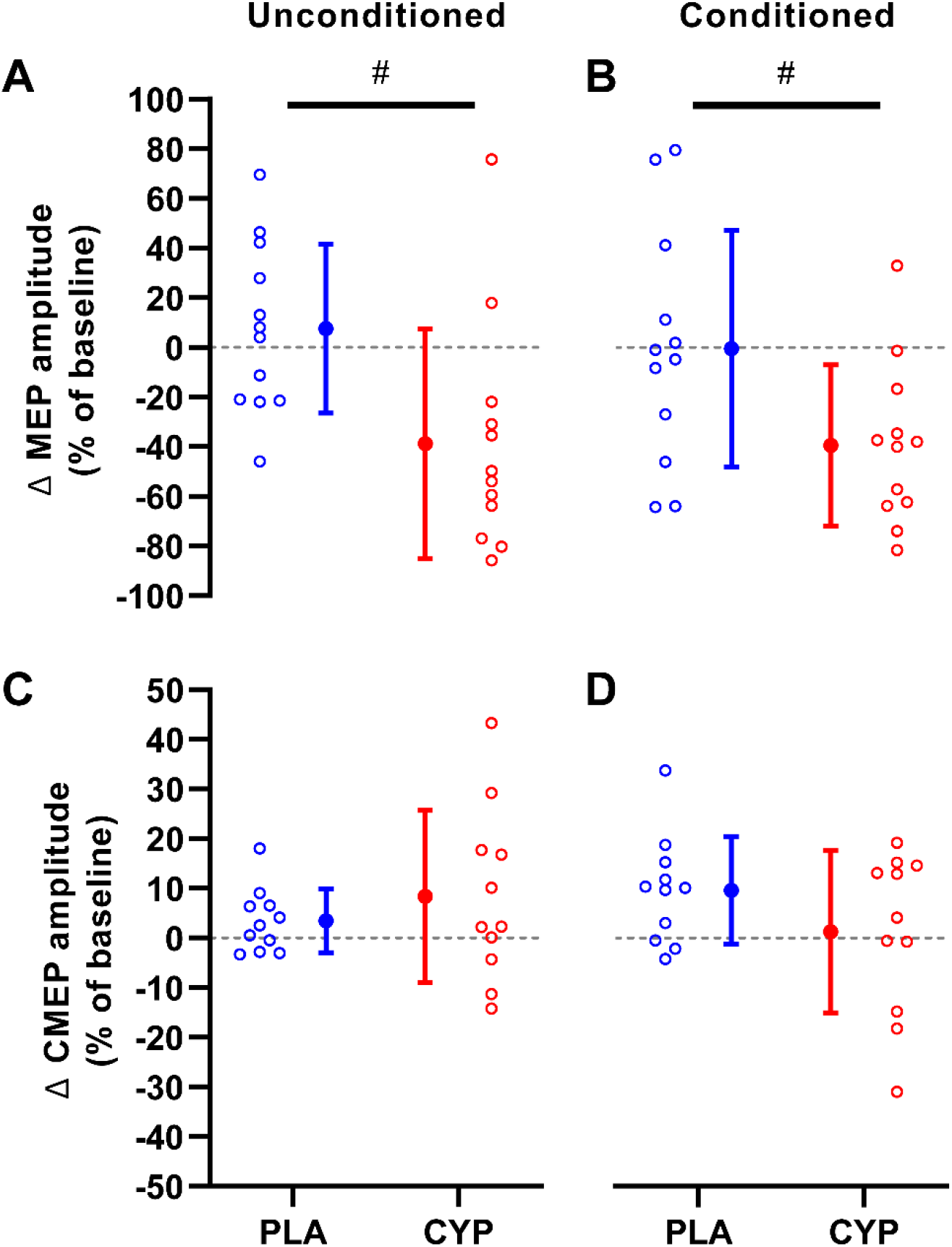
MEP and CMEP changes with 5-HT_2_ receptor blockade while the muscle was at rest. Unconditioned MEP (A), conditioned MEP (B), unconditioned CMEP (C), and conditioned CMEP (D) were measured before and after the ingestion of a placebo or cyproheptadine. Differences between pre-pill data and post-pill data were normalised to pre-pill data, where a negative value indicates that the post-pill value was lower than the pre-pill value. Data are presented as the group mean and standard deviation (MEPs: n = 12, CMEPs: n = 11). Individual data are presented for the placebo session (blue circles) and the cyproheptadine session (red circles). Hash symbols represent a significant difference between placebo and cyproheptadine data (p < 0.05).

**Figure 5.**
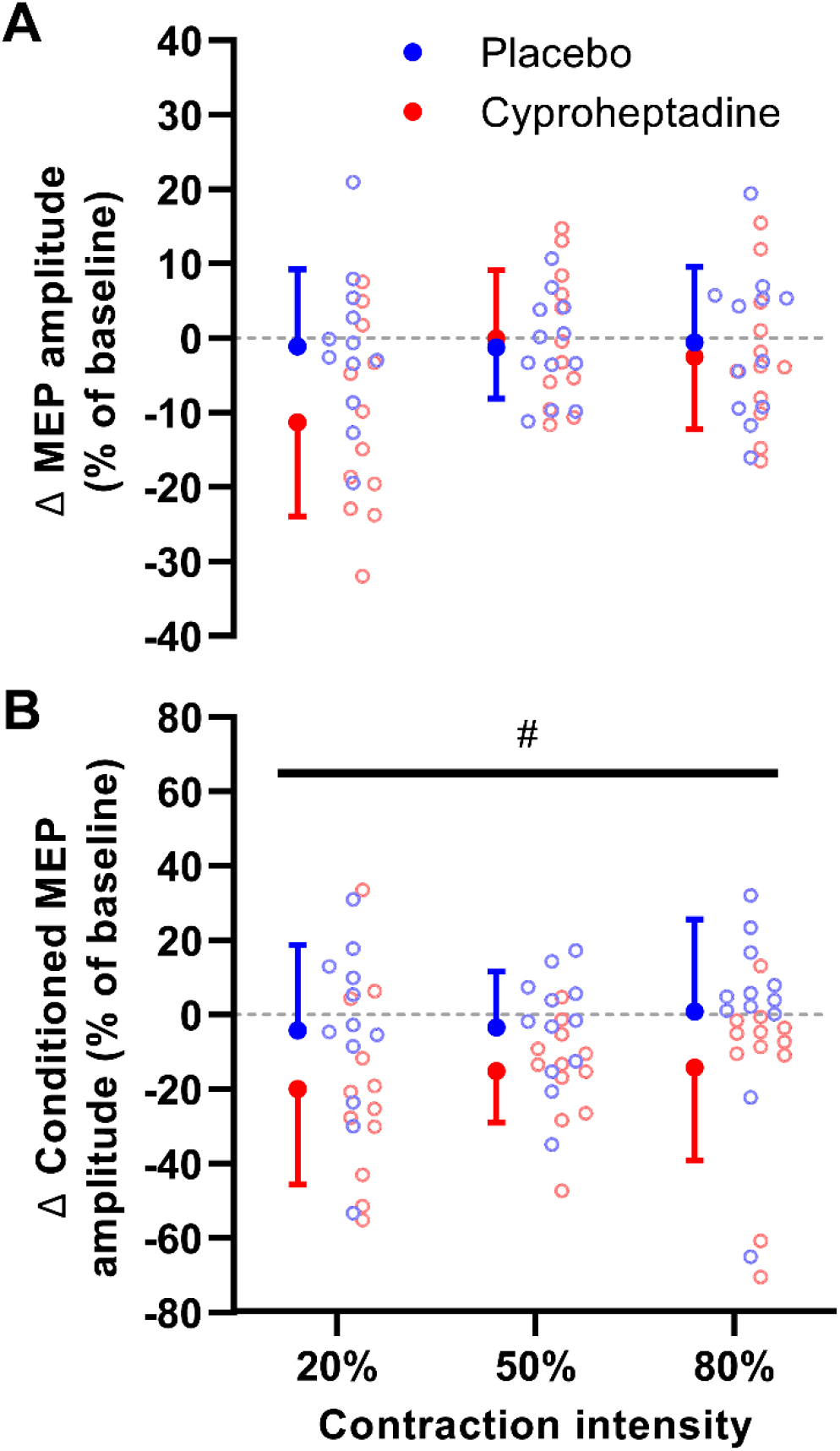
MEP changes with 5-HT_2_ receptor blockade during submaximal contractions. TMS-evoked MEPs were measured from the biceps brachii during the elbow flexions of 20% MVC, 50% MVC, and 80% MVC (A). MEPs were also obtained during these submaximal contractions when a TMS conditioning pulse was delivered 100 ms prior to the test pulse (B). Post-pill data were normalised to pre-pill data, where a negative value indicates that the post-pill value was lower than pre-pill. Data are presented as the group mean and standard deviation (n = 12). Individual data are presented for the placebo session (blue circles) and the cyproheptadine session (red circles). Hash symbols represent a significant main effect of drug (p < 0.05).

**Table 3.**
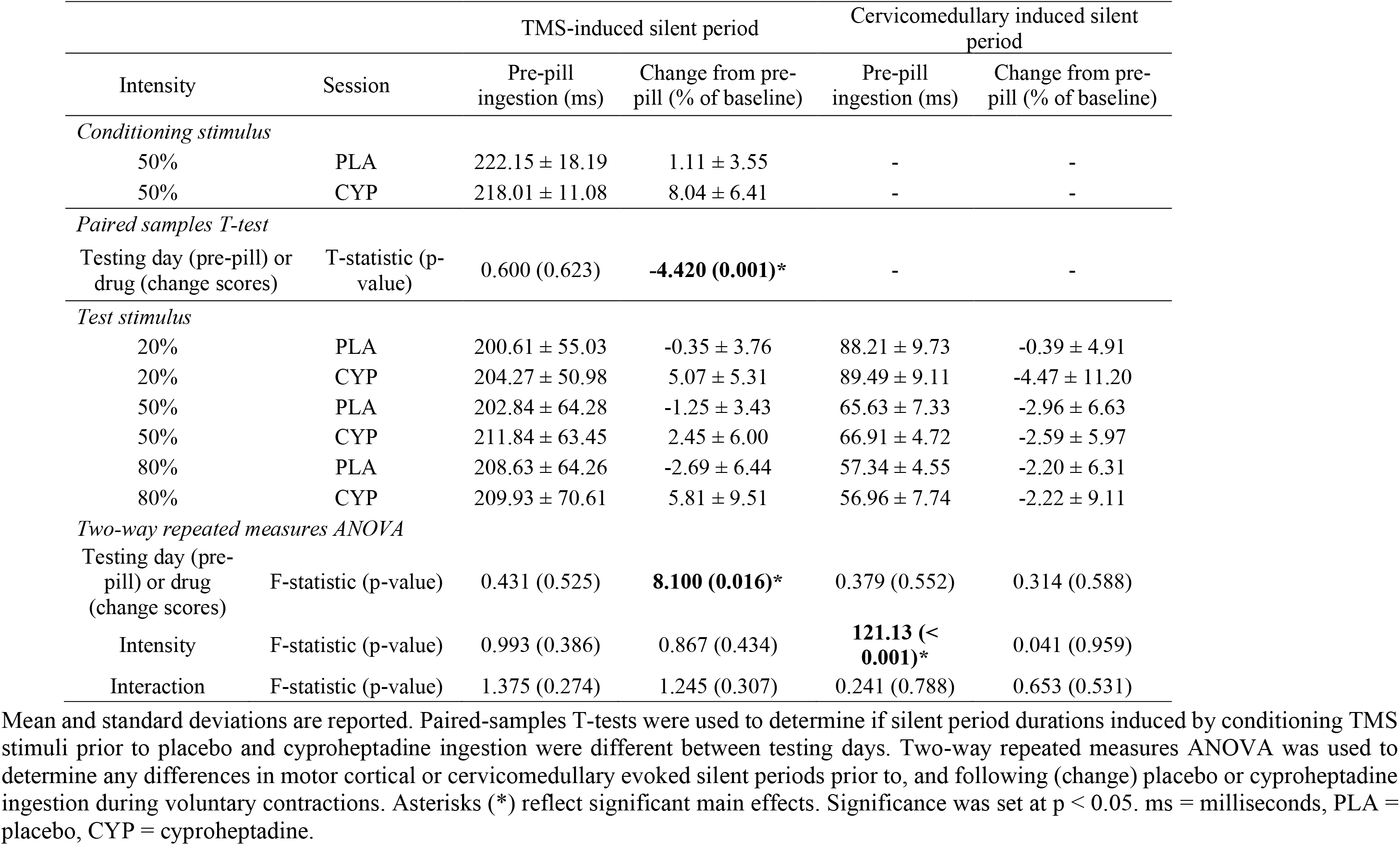
Table of silent period durations induced by motor cortical (TMS) and electrical cervicomedullary stimulation.

|  |  | TMS-induced silent period |  | Cervicomedullary induced silent period |  |
| --- | --- | --- | --- | --- | --- |
| Intensity | Session | Pre-pill ingestion (ms) | Change from pre-pill (% of baseline) | Pre-pill ingestion (ms) | Change from pre-pill (% of baseline) |
| *Conditioning stimulus* |  |  |  |  |  |
| 50% | PLA | 222.15 ± 18.19 | 1.11 ± 3.55 | - | - |
| 50% | CYP | 218.01 ± 11.08 | 8.04 ± 6.41 | - | - |
| *Paired samples T-test* |  |  |  |  |  |
| Testing day (pre-pill) or drug (change scores) | T-statistic (p-value) | 0.600 (0.623) | **-4.420 (0.001)\*** | - | - |
| *Test stimulus* |  |  |  |  |  |
| 20% | PLA | 200.61 ± 55.03 | -0.35 ± 3.76 | 88.21 ± 9.73 | -0.39 ± 4.91 |
| 20% | CYP | 204.27 ± 50.98 | 5.07 ± 5.31 | 89.49 ± 9.11 | -4.47 ± 11.20 |
| 50% | PLA | 202.84 ± 64.28 | -1.25 ± 3.43 | 65.63 ± 7.33 | -2.96 ± 6.63 |
| 50% | CYP | 211.84 ± 63.45 | 2.45 ± 6.00 | 66.91 ± 4.72 | -2.59 ± 5.97 |
| 80% | PLA | 208.63 ± 64.26 | -2.69 ± 6.44 | 57.34 ± 4.55 | -2.20 ± 6.31 |
| 80% | CYP | 209.93 ± 70.61 | 5.81 ± 9.51 | 56.96 ± 7.74 | -2.22 ± 9.11 |
| *Two-way repeated measures ANOVA* |  |  |  |  |  |
| Testing day (pre-pill) or drug (change scores) | F-statistic (p-value) | 0.431 (0.525) | **8.100 (0.016)\*** | 0.379 (0.552) | 0.314 (0.588) |
| Intensity | F-statistic (p-value) | 0.993 (0.386) | 0.867 (0.434) | **121.13 (< 0.001)\*** | 0.041 (0.959) |
| Interaction | F-statistic (p-value) | 1.375 (0.274) | 1.245 (0.307) | 0.241 (0.788) | 0.653 (0.531) |

Mean and standard deviations are reported. Paired-samples T-tests were used to determine if silent period durations induced by conditioning TMS stimuli prior to placebo and cyproheptadine ingestion were different between testing days. Two-way repeated measures ANOVA was used to determine any differences in motor cortical or cervicomedullary evoked silent periods prior to, and following (change) placebo or cyproheptadine ingestion during voluntary contractions. Asterisks (\*) reflect significant main effects. Significance was set at $p < 0.05$ . ms = milliseconds, PLA = placebo, CYP = cyproheptadine.

Overall, changes in CMEP amplitude due to 5-HT_2_ receptor blockade were the opposite of MEP amplitude. Drug-effects were detected for the unconditioned CMEP, where cyproheptadine caused a significantly greater reduction in unconditioned CMEP amplitude compared to placebo (main effect of drug: F(1, 10) = 5.493, p = 0.041, η^2^ = 0.355, Figure 6A). Changes in the unconditioned CMEP were not dependent on the intensity of the elbow flexion, as there was no main effect of contraction intensity (p = 0.924), and no drug by contraction intensity interaction (p = 0. 919) identified for unconditioned CMEP amplitude. There were no significant changes in conditioned CMEP amplitude due to cyproheptadine or the submaximal contraction being performed, as there was no main effect of drug (p = 0.667), contraction intensity (p = 0.562), or drug by contraction intensity interaction (p = 0.962) detected for conditioned CMEPs (Figure 6B). There were no differences to the silent period duration induced by singe cervicomedullary stimulation following cyproheptadine ingestion (see Table 3).

**Figure 6.**
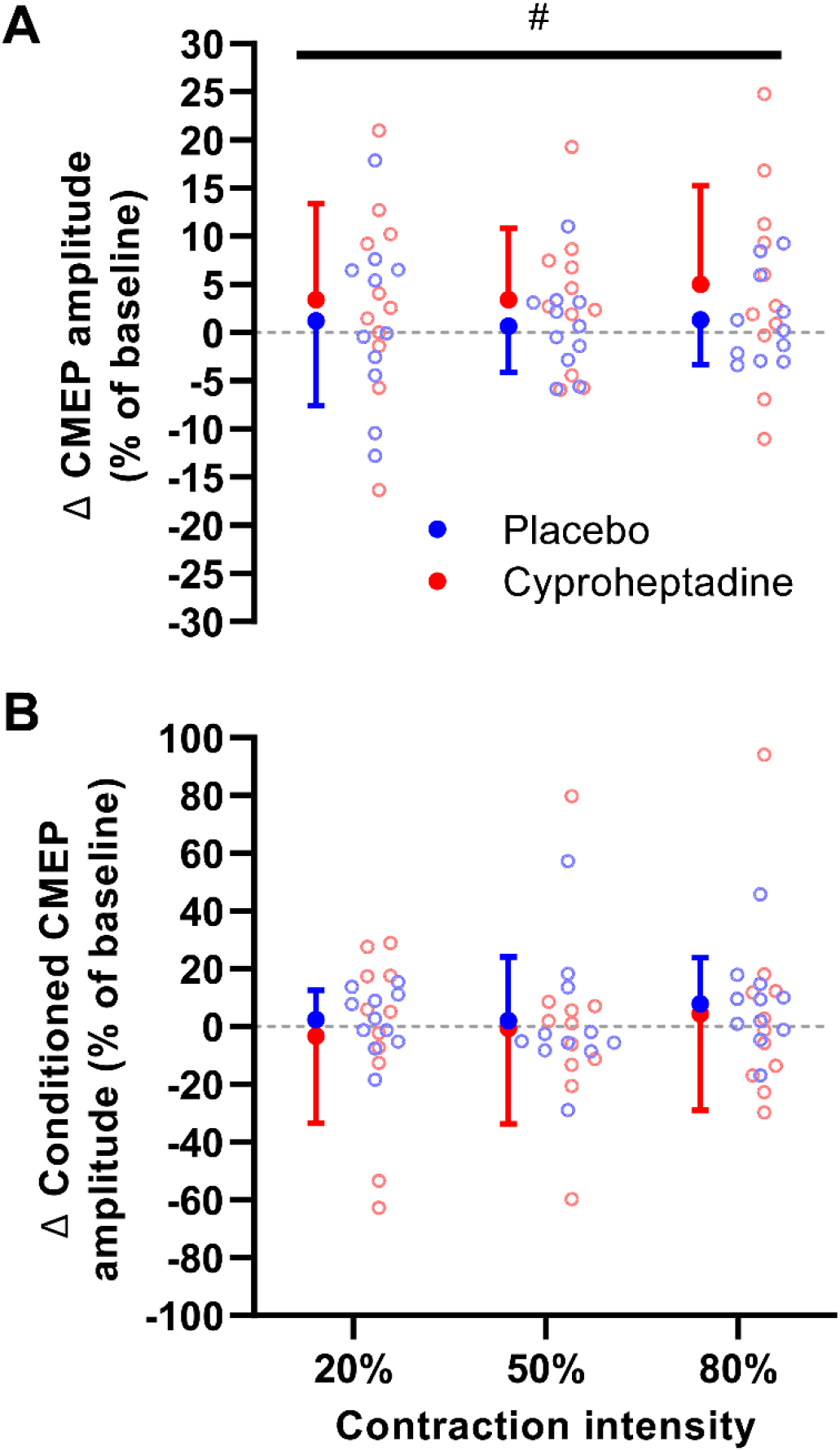
CMEP changes with 5-HT_2_ receptor blockade during submaximal contractions. CMEPs were measured from the biceps brachii during the elbow flexions of 20% MVC, 50% MVC, and 80% MVC (A). CMEPs were also obtained during these submaximal contractions when a TMS conditioning pulse was delivered 100 ms prior to the test pulse (B). Post-pill data were normalised to pre-pill data, where a negative value indicates that the post-pill value was lower than pre-pill. Data are presented as the group mean and standard deviation (n = 11). Individual data are presented for the placebo session (blue circles) and the cyproheptadine session (red circles). Hash symbols represent a significant main effect of drug (p < 0.05).

Not surprisingly, the RMS amplitude of biceps brachii EMG was significantly different for the contraction intensity (F(1, 11) = 186.338, p < 0.001, η^2^ = 0.944), where EMG RMS amplitude progressive increased from 20% MVC to 50% MVC to 80% MVC (Table 2). The change in EMG amplitude from pre-pill to post-pill was unaffected by the drug. Given that the change in M_MAX_ from pre-pill to post-pill was also unaffected by the drug it is likely that drug-effects were associated with the CNS rather than the muscle during submaximal contractions.

## DISCUSSION

This study investigated how 5-HT_2_ receptor antagonism affected corticospinal and motoneuronal excitability in humans. MEPs and CMEPs were obtained from the biceps brachii with, and without, a preceding TMS conditioning stimulus. This allowed us to examine the effects of 5-HT_2_ receptor antagonism on MEPs and CMEPs when descending drive was being delivered to the muscle during voluntary muscle contractions as well as during a period of motoneurone disfacilitation. The main findings of this study were that 5-HT_2_ receptor antagonism caused 1) a reduction in conditioned and unconditioned MEP amplitude when the muscle was at rest, 2) a reduction in conditioned MEPs across a range of submaximal contractions, and 3) an increase in unconditioned CMEP amplitude across a range of submaximal contractions. In addition, 5-HT_2_ antagonism caused a reduction in MVCs which confirms previous findings. Collectively, these novel findings suggest 5-HT_2_ receptors modulate intracortical excitability and motoneuronal excitability differently.

### Unconditioned and conditioned MEPs from the biceps brachii

A single TMS stimulus applied to the motor cortex generates multiple descending volleys to the motoneurone pool to produce an excitatory MEP response in the EMG signal (Hupfeld et al., 2020; Kojima et al., 2013; Zeugin & Ionta, 2021). The size of the unconditioned MEP is dependent on the stimulus intensity as well as the level of voluntary effort (Rothwell et al., 1987; Taylor et al., 1997). In the current study there was a clear difference in unconditioned MEP amplitude for MEPs obtained at rest compared to MEPs obtained during voluntary contractions. In the relaxed muscle, the MEP size is particularly sensitive to stimulus intensity, where a change in TMS intensity from resting motor threshold (RMT) to 160% RMT can increase unconditioned MEP size by almost 20-fold (McNeil, Martin, et al., 2011). However, the sensitivity to TMS stimulus intensity is reduced during voluntary contractions, which is particularly obvious at higher forces due to motoneurones reducing their responsiveness to the synaptic input when firing at high rates (Martin et al., 2006). The large difference between unconditioned MEP size at rest compared to unconditioned MEPs during contractions is most likely due to increased excitability of the motor cortex as well as the motoneurone pool (Di Lazzaro et al., 1998; McNeil et al., 2013).

MEPs were also produced during the period of EMG silence that was generated by a conditioning TMS pulse. This paired-stimulus technique likely has effects at two levels. It is suggested to reduce the influence of muscle contraction on conditioned responses by interrupting descending voluntary drive and placing the motoneurone pool in a similar state to rest (Fuhr et al., 1991; McNeil et al., 2009; McNeil, Martin, et al., 2011). In addition, when MEPs are obtained 100 ms following a conditioning stimulus, the amplitude of the conditioned MEP reflects long-interval intracortical inhibition (LICI) (Florian et al., 2008; Valls-Sole et al., 1992). The cortical component of this paired-pulse technique is suggested to reflect inhibition mediated by γ-amino butyric acid (GABA_b_) receptors located throughout intracortical circuits (Florian et al., 2008; McDonnell et al., 2006). GABA_b_ receptors are G-protein coupled and modulate potassium and calcium ion channels (Nicoll, 2004) to promote long-lasting inhibitory post-synaptic potentials (IPSPs) in motor cortical output cells (Florian et al., 2008; McDonnell et al., 2006).

### Intracortical inhibition is enhanced with 5-HT_2_ antagonism

5-HT_2_ receptor antagonism reduced unconditioned and conditioned MEPs at rest, and only affected conditioned MEP amplitude during contractions. Therefore, it appears that 5-HT_2_ receptor antagonism only influenced motor cortical circuits when there was no muscle activity or when LICI was evoked. We propose that the changes to MEPs and LICI with 5-HT_2_ antagonism are primarily due to modulation of intracortical circuits, given that motoneurones are close to a resting state when a conditioned MEP is evoked, and 5-HT_2_ antagonism does not affect CMEPs at rest (Thorstensen et al., 2022). A lack of differences for resting CMEPs following ingestion of a 5-HT_2_ antagonist indicate that there is possibly no change to the responsiveness of the motoneurone pool to synaptic input at rest, so differences in MEPs are probably due to changes in supraspinal circuits. 5-HT_2_ receptors located on GABA neurons can influence GABA release (Ciranna, 2006; Shen & Andrade, 1998) and the generation of GABA_b_ IPSPs (Oleskevich & Lacaille, 1992). However, 5-HT_2_ antagonism reduced the amplitude of conditioned MEPs, suggesting stronger intracortical inhibition when 5-HT_2_ receptors are less active. Nonetheless, the observed increase in intracortical inhibition with 5-HT_2_ antagonism is possibly due to an interaction between 5-HT_2_ and GABA_b_ receptors.

This is not the only study to propose an interaction between the serotonergic system and TMS measures of GABAergic activity. Several recent human experiments have revealed a 5-HT mediated role in cortical inhibition, whereby ingestion of a selective serotonin reuptake inhibitor (SSRI) can lengthen the TMS-induced silent period (Henderson et al., 2022; Robol et al., 2004; Thorstensen et al., 2020) and increase short-interval intracortical inhibition (SICI) associated with GABA_a_ receptors (Robol et al., 2004). Given that SSRIs increase SICI, and 5-HT_2_ antagonism increases LICI, it is probable that a serotonergic mechanism influences cortical inhibition. As both GABA_a_ and GABA_b_ receptors have been implicated it is likely that no single neuromodulatory mechanism is at play. Nonetheless, the findings of our study provide evidence that 5-HT_2_ receptors can influence corticospinal excitability through modulation of intracortical inhibition. However, since 5-HT_2_ antagonism did not alter MEPs when descending drive was being delivered to the spinal cord, 5-HT_2_ effects in motor cortical circuits do not appear to influence corticospinal excitability during submaximal voluntary contractions.

### Unconditioned and conditioned CMEPs from the biceps brachii

Electrical stimulation was delivered to the cervicomedullary junction in the current study to measure spinal motoneurone responsiveness to synaptic input. Cervicomedullary stimulation generates a single descending volley through excitation of corticospinal axons to activate motoneurones and produce a CMEP as a measure of motoneurone excitability (McNeil et al., 2013; Taylor, 2006; Taylor & Gandevia, 2004). Unconditioned CMEPs increased in size from rest to contractions, but at a reduced magnitude of change compared to the unconditioned MEP. Unlike motor cortical stimulation, subcortical electrical stimuli applied to the cervicomedullary region directly excites corticospinal axons at the pyramidal decussation (Petersen et al., 2002; Taylor, 2006; Ugawa et al., 1991; Yacyshyn et al., 2016). For biceps brachii, cervicomedullary stimulation often activates a larger population of motoneurones at rest compared to motor cortical TMS, with a large direct/monosynaptic component to the response. The increase in size from rest to voluntary efforts in the current study support that motoneurone facilitation occurs as descending drive to the motoneurone pool brings motoneurones closer to firing thresholds to become more responsive to the consistent synaptic input provided by electrical corticospinal axon stimuli.

The current study revealed that both conditioned MEPs and CMEPs are reduced during voluntary contractions compared to unconditioned responses. However, conditioned MEPs linearly increased with contraction intensity whereas conditioned CMEPs linearly decreased with contraction intensity. Our findings for conditioned MEPs and contraction intensity align with previously reported LICI protocols employing voluntary contractions (McNeil, Martin, et al., 2011). However, changes to conditioned CMEP size during a broad range of submaximal voluntary contraction have not been previously reported. Conditioned CMEPs were typically smaller than unconditioned CMEPs, which is presumably due to motoneurone disfacilitation, and may indicate a spinal contribution to LICI (McNeil, Giesebrecht, et al., 2011; McNeil, Martin, et al., 2011; Yacyshyn et al., 2016). Disfacilitation of motoneurones would place the motoneurone pool in a similar state to rest, but since the conditioned CMEP was smaller during the contractions compared to the unconditioned CMEP at rest, motoneurones were probably more difficult to recruit during the TMS-induced silent period compared to a relaxed muscle. Thus, it appears that motoneurones were not only disfacilitated, but also inhibited in the TMS-induced silent period. The magnitude of inhibition was also larger during stronger contractions. Interestingly, we observed larger conditioned MEPs than conditioned CMEPs, which was most evident during high intensity contractions (80% MVC). Thus, the motor cortical stimulation was able to activate more of the motoneurone pool than cervicomedullary stimulation during the silent period.

### 5-HT2 antagonism exerts effects on spinal motoneurones only when descending drive is present

Both the unconditioned CMEP and the conditioned CMEP were unaffected by 5-HT_2_ antagonism in the resting muscle, and there were no 5-HT related changes to conditioned CMEP size during submaximal contractions. However, 5-HT_2_ antagonism increased CMEP amplitude during the submaximal contractions when descending drive was still being delivered to the spinal cord. Given 5-HT_2_ modulation of motoneurone excitability is primarily mediated by dendritic PICs (Perrier & Delgado-Lezama, 2005; Perrier & Hounsgaard, 2003), and PICs require sustained excitatory inputs to become active (Heckman et al., 2008), voluntary drive is probably needed to reveal 5-HT modulation via PICs. Our previous work demonstrated that 5-HT_2_ antagonism does not affect (unconditioned) CMEP amplitude in the resting biceps brachii, even when voluntary contractions were performed in the opposite limb to enhance 5-HT concentration in the spinal cord (Thorstensen et al., 2022). These previous results, and the results of the current study are consistent with the need for sustained depolarizing input to activate PICs on the motoneurone, and that reduced PIC facilitation with 5-HT_2_ antagonism will not reduce motoneurone responsiveness to a brief synaptic volley where there is no background activation of motoneurones.

The increased CMEP with 5-HT_2_ antagonism suggests that when a cervicomedullary stimulus was delivered, more motoneurones were responsive to the additional synaptic excitatory post-synaptic potential, and more motoneurones were recruited into the evoked response. This could be explained by differences in motoneurone firing rates and recruitment with 5-HT_2_ antagonism. Indeed, recent human investigations using lower limb high-density surface EMG report a reduction to motor unit discharge rate following ingestion of a 5-HT_2_ antagonist (Goodlich et al., 2022). Such changes could be brought about by a change in the balance of synaptic inputs driving the motoneurone pool. Consistent with this idea, 5-HT_2_ antagonism was found to reduce estimates of PICs during low intensity dorsiflexion contractions (Goodlich et al., 2023). This same effect may be present in the current study as 5-HT_2_ antagonism also reduced maximal elbow flexion torque. For submaximal contractions, this would mean that extra voluntary drive to the motoneurones would be required to compensate for the reduced PICs, and could cause increased firing rates and/or recruitment of motor units. Moreover, given that post-pill submaximal targets were based on pre-pill MVC, submaximal torque targets were relatively larger with 5-HT_2_ antagonism. Thus, voluntary drive to motoneurones must be stronger to perform the submaximal contraction with 5-HT_2_ antagonism, and more motoneurones would be 1) closer to firing threshold (i.e., in the subliminal fringe), and 2) more readily recruited into the CMEP. A further consideration is that when PICs are active the responsiveness to additional synaptic inputs to the motoneurone is limited (Heckman et al., 2008). Reducing the action of PICs via 5-HT_2_ antagonism may allow the motoneurone to retain its responsiveness to the synaptic inputs.

## Conclusion

This study provides novel insight to the complex serotonergic modulation of motoneurone excitability in humans. Antagonism of 5-HT_2_ receptors within the CNS increased intracortical inhibition and increased CMEP amplitude during voluntary efforts. This is the first study to reveal 5-HT_2_ receptor involvement in LICI, suggesting a 5-HT_2_ and GABAergic interaction within motor cortical pathways. The results also indicate that excitatory drive to the spinal cord is necessary to observe the effects of 5-HT_2_ antagonism on spinal motoneurone excitability.

